# Genomic and transcriptomic profiling of high-risk bladder cancer reveals diverse molecular and microenvironment ecosystems

**DOI:** 10.1101/2024.12.21.629010

**Authors:** Khyati Meghani, Yanni Yu, Noah Frydenlund, Erik Z. Li, Bonnie Choy, Sarki A. Abdulkadir, Joshua J. Meeks

**Author notes:** Corresponding author, Joshua J. Meeks,; @JoshMeeks, C (312)363-8959.

## Abstract

Despite surgical resection, rigorous endoscopic surveillance, and immunotherapy with the Bacillus Calmette–Guérin (BCG) vaccine, 30% of high-risk bladder cancers recur, and 10% result in fatal outcomes within two years of diagnosis. The global shortage of BCG underscores the urgent need for alternative or complementary therapeutic strategies. To address this, we integrated transcriptomic profiling and targeted genomic sequencing to identify four consensus intrinsic subtypes of bladder cancer. Initially derived from bulk RNA profiling, these subtypes were further validated at the cellular and tissue-compartment levels using single-cell RNA sequencing and spatial transcriptomics. Notably, we identified a subtype of inflamed tumors with enhanced endogenous retroelement expression and increased commensal bacterial presence, which showed the highest responsiveness to BCG therapy. Additionally, we developed a machine learning-based model incorporating composite molecular features to predict recurrence risk, achieving a high accuracy (AUC = 0.90). Our findings establish a molecular precision framework for bladder cancer and nominate novel therapeutic targets to reduce reliance on BCG immunotherapy.

## Introduction

Nearly 80% of bladder cancer diagnoses are non-muscle invasive (NMIBC, Stage T1 or less) treated with surgical resection and intracavitary treatment with immunotherapy for 36 months^1^. Yet, 26% of T1 tumors will recur within two years, and 10% will progress to metastasis and will be fatal. Bacillus Calmette Guérin (BCG), the tuberculosis vaccine, is the most effective agent to decrease recurrence and progression and has been the primary oncologic treatment of T1 cancers for over 50 years^2^. However, production shortages have produced a global scarcity of BCG, with no foreseeable improvement in the near future^3^. Consequently, there is an urgent need to identify alternative therapeutic targets that can effectively augment or replace BCG in patients with limited access or poor response^4^.

Despite its long-term therapeutic benefit in bladder cancer, the mechanisms of response to BCG therapy are not known. Most T1 cancers have limited immune infiltration, and the molecular features of the tumor affecting the tumor microenvironment (TME) remain unknown^5^. A complex interaction between the tumor and autocrine/paracrine immune regulatory circuits within the TME is hypothesized to mediate the immune response to BCG. Mechanistically, it is unclear if the immune response resulting in recurrence-free survival is anti-BCG (a tuberculosis-specific response), anti-tumor, or both^6–11^. Enrichment of the TME with CD4+ T cells^7^ is associated with an improved response to BCG therapy. Furthermore, exhaustion of these CD4+ T cells, with elevated PD-L1 expression, predicts a lack of response to BCG^12^. Microbial populations that inhabit the mucosal surfaces of the bladder microenvironment may also play a role in transforming the underlying epithelium into an “inflamed” state and augmenting the immune response^13^. Empiric research has identified a relationship between the gut microbiome and anti-tumor immunity in bladder cancer^14–19^. We and several others have previously investigated transcriptomic signatures of BCG response for T1 stage cancers^6,20,21^. However, despite multiple attempts to classify tumors based on transcriptomic subtypes, a comprehensive atlas of actionable characteristics encompassing the observed heterogeneity in T1 NMIBCs is lacking.

In this study, we identify four distinct molecular subtypes based on expression profiling of T1 tumors characterized by unique features of transcriptomic and genomic profiles. Employing phylogenetic analysis, we uncovered microbial species within the bladder TME of one subtype associated with elevated pre-treatment immune infiltration and a more prolonged recurrence-free survival after BCG therapy. Using machine learning, we extract RNA single-base-substitution signatures revealing ongoing APOBEC activity in tumors. By integrating these diverse clinical and molecular features, we identify molecular subtypes exhibiting distinctive biology that nominate precision therapeutic interventions for high-risk bladder cancer

## Results

### Unsupervised clustering approach to identify drivers of underlying heterogeneity in T1 non-muscle invasive bladder cancer

We performed transcriptional profiling of 106 high-risk (Stage T1) tumors with targeted mutational sequencing of 92 tumors using a bespoke 194-gene panel designed for T1-stage bladder cancers. We identified four tumor subtypes by unsupervised clustering of normalized expression values, focusing on the 1,500 genes with the highest variation **(Supp Fig 1A**). Comparing these clusters to those previously identified by Robertson et al.^20^, the only other stage-specific profiling, we calculated an adjusted mutual information score of only 0.29, suggesting distinct differences in the expression profiles captured by each clustering approach **(Supp Fig 1B**).

**Figure 1:**
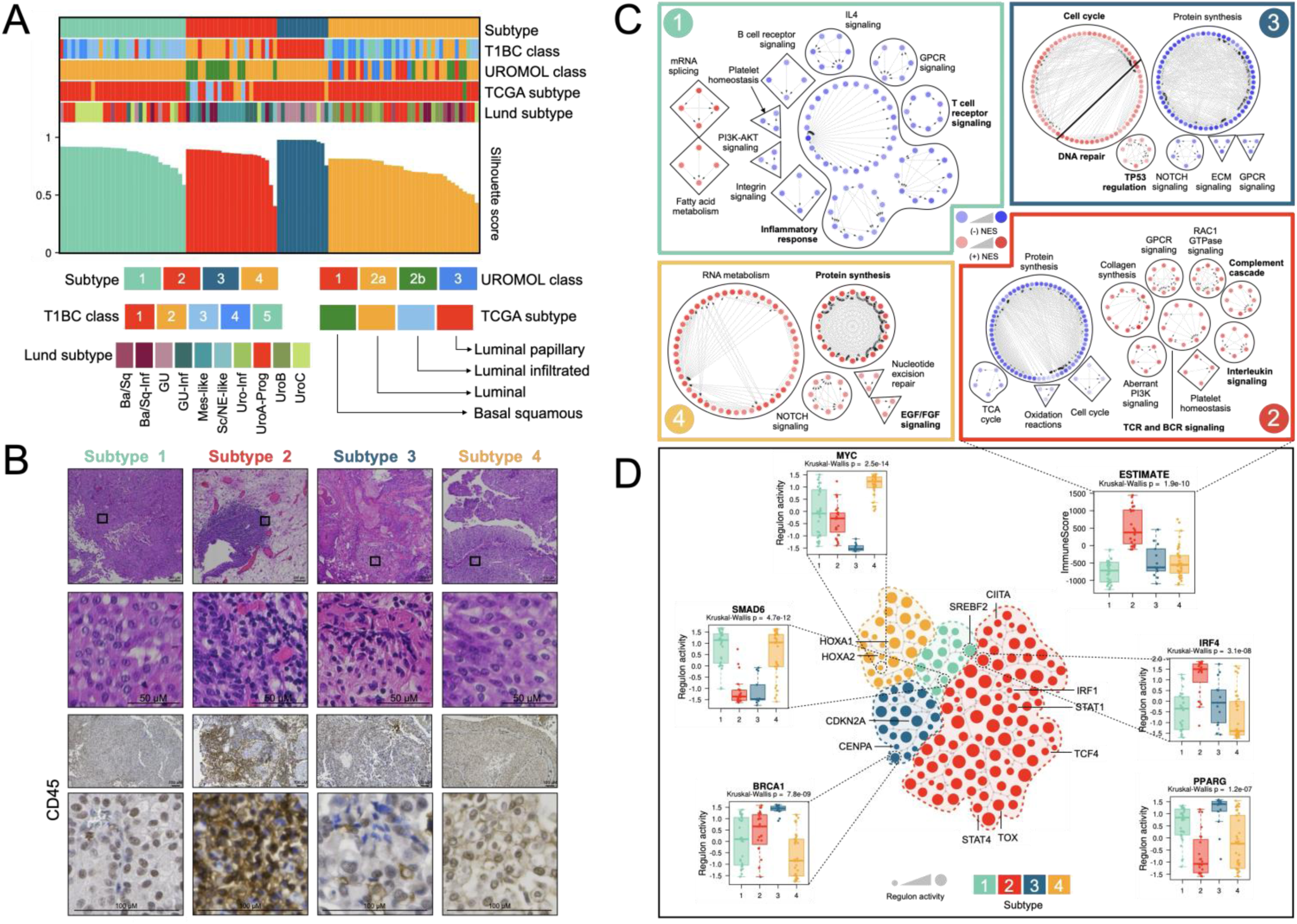
Unsupervised clustering reveals four molecular groups of high-risk bladder cancers. A. Covariate tracks displaying the predicted subtypes of 106 T1 tumors using the T1BC, UROMOL, TCGA and Lund classifiers. B. Representative H&E images for tumor within each Subtype. (Top panel). Representative immunohistochemistry images showing CD45 staining in tumors. (Bottom panel). C. Distinguishing gene program networks in each Subtype visualized as a Cytoscape network. Each node represents a pathway and edges represent shared genes between the connecting pathways. Nodes colored in red are upregulated and nodes colored in blue are downregulated gene sets in each Subtype group. D. Central tree-and-leaf plot shows the unique regulons identified in each Subtype. Surrounding boxplots show regulon activity profiles for highlighted genes and Immune score by Subtype. (p-values were calculated from a two-sided Kruskal-Wallis test)

Classifying the tumors in our cohort using MIBC TCGA (Th Cancer Genome Atlas) expression classification^22^, we find most tumors were classified as luminal papillary (93/106, 88%) (Fisher’s Exact test p-value = 3.29e-05). The remaining tumors were distributed among luminal infiltrated (6/106, 5.6%), luminal (5/106, 4.7%), with few basal squamous (2/106, 1.8%) tumors. We applied the UROMOL classifier^6^ and found 66% of tumors classified as Class 2a (70/106), 17% as Class 2b (18/106), 9.4% as Class 1 (10/106), and 7.5% as Class 3 (8/106) (**Fig 1A**). Our findings align with those reported by the UROMOL cohort in which Class 2a was enriched for Stage I tumors^6^. Classifying tumors using the Lund subtyping method^23,24^, we find 78% of Subtype1 tumors (25/32) and 76% of Subtype4 (29/38) tumors fall within the Urothelial-like (Uro) subtype comprising of Uro-Inf, UroA-Prog, UroB and UroC. 92% of Subtype3 tumors (12/13) are classified within the genomically unstable (GU) subtypes (GU and GU-Inf). 47% of Subtype2 tumors (11/23) were classified as Mesenchymal-like (Mes-like), 21% (5/23) were classified as basal-squamous like (Ba/Sq-Inf), and another 21% (5/23) were classified as GU-Inf.

We then evaluated the clinical and histologic features of each subtype. We first performed a histologic comparison (**Fig 1B**, top panel). Subtypes 1 and 4 had papillary architecture, and tumor cells from Subtype3 had larger, pleomorphic nuclei. By comparison, Subtype2 tumors infiltrated with immune cells (CD45+) (**Fig 1B**, bottom panel). Next, we analyzed recurrence-free survival among tumor subtypes after BCG treatment to determine if the molecular differences affected response to BCG. We observed that three of the four transcriptomic subtypes had poor outcomes: 53% of Subtype1 tumors (17/32 recurred at 24 months, median survival 43.6 months), 50% of Subtype3 (6/12 recurred at 24 months, median survival 82 months), and 62% of Subtype4 (16/37 recurred at 24 months, median survival 34.1 months) tumors had a recurrence at 24-months. One subtype (Subtype2) demonstrated an improved recurrence-free survival : 68.2% recurrence-free (15/22) at 24 months **(Supp Fig 1C**; Log-rank p = 0.023).

### Transcriptomic diversity in NMIBC

We investigated the gene expression programs associated with each Subtype (**Fig 2A**). Subtype1 tumors expressed downregulated immune signatures with enrichment of mRNA splicing and fatty acid metabolism pathways. In contrast, Subtype2 tumors were enriched with immune, interleukin, and cytokine signaling pathways, accompanied by a higher RNA-based Immune score by ESTIMATE analysis^25^. These findings align with the enhanced immune infiltration in Subtype2 tumors highlighted by CD45+ immunohistochemistry (**Fig 1B**). Subtype3 tumors were enriched in pathways related to cell-cycle regulation and DNA damage response and repair. Finally, FGF signaling, protein synthesis, and RNA metabolism pathways were enhanced in Subtype4 tumors w (**Fig 1C, Fig 7**). To further dissect the transcriptional regulatory networks driving these gene programs, we explored the regulon networks^22,26^ within each Subtype (**Fig 1D**). Elevated regulon activities of SREBF2, a regulator of fatty acid synthesis, SMAD6, genes associated with immune exclusion and anti-inflammation^27^ were identified in Subtype1 tumors. Consistent with the pathway analysis, we identified several upregulated inflammatory regulons within the STAT and IRF protein families in Subtype2. A striking feature of Subtype2 tumors was the significant downregulation of the PPARG regulon observed in this subtype. Elevated PPARG expression has been shown to drive immune exclusion in bladder tumors^28^, and the downregulation of the PPARG regulatory network within Subtype2 could be a key driver of the observed inflamed phenotype in these tumors.

**Figure 2:**
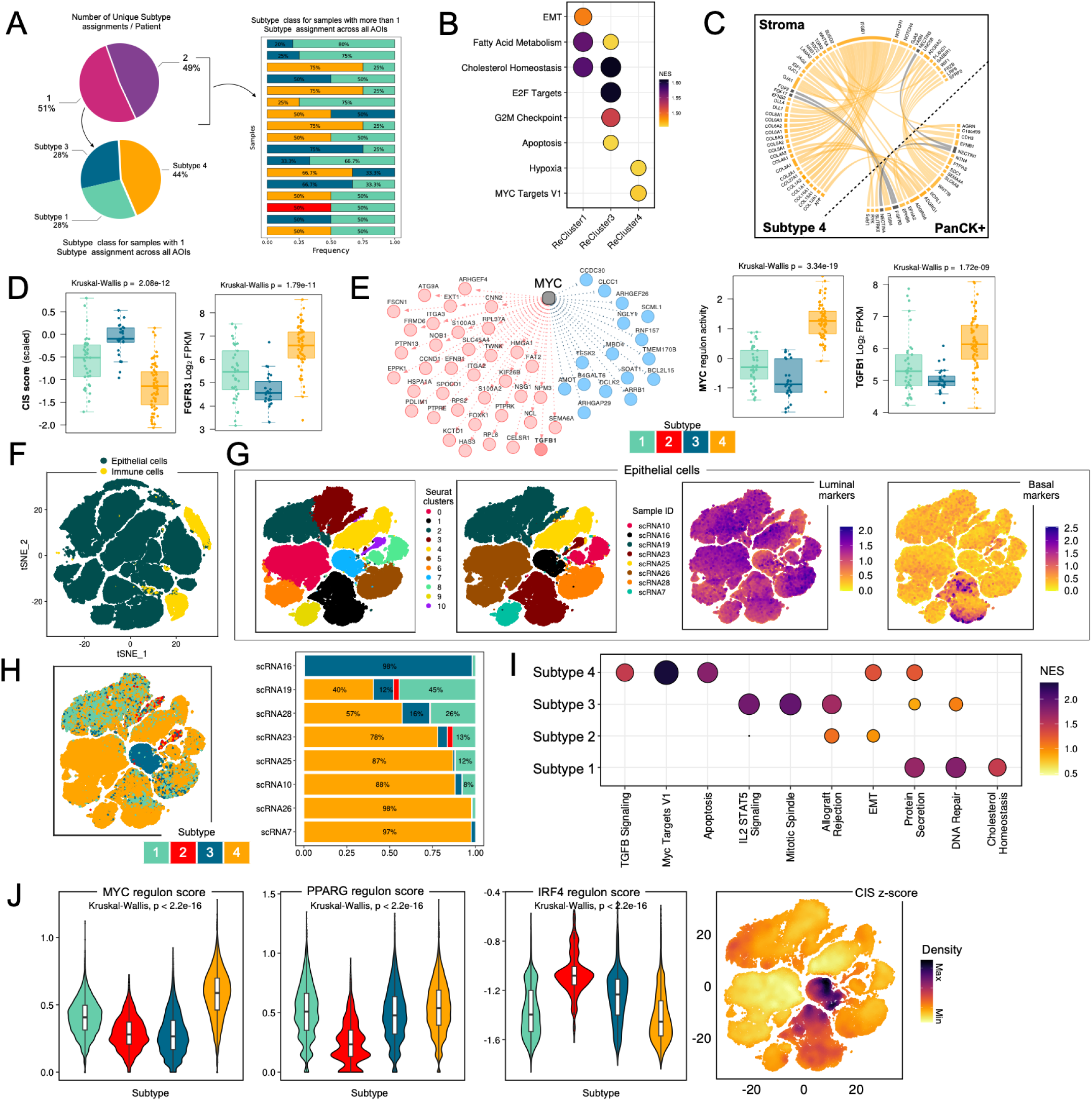
Heterogeneity in identified programs dissected using spatial and single cell transcriptomic profiling. A. Pie-chart showing the frequency of intra-tumor heterogeneity in Subtype expression in a digital spatial profiling dataset of non-muscle invasive tumors (Top, left). Pie chart showing the frequency of Subtype classification for patients with homogenous Subtype classification across all AOIs. (Bottom, left). Bar chart showing the Subtype assignments for patients with greater than 1 Subtype assignment across all AOIs. (Right). B. Dot plot showing significant gene programs differentially expressed in PanCK+ AOIs assigned to each Subtype. C. Circos plot showing strength of interaction between highlighted cytokine-receptor pairs in adjacent PanCK and Stroma AOI. D. Boxplot showing CIS score (left) and FGFR3 mRNA levels (right) across PanCK+ AOIs colored by Subtype class. E. Left: MYC regulon network highlighting activating targets (red) and inhibitory targets (blue). Right: Boxplot showing levels of MYC activating regulon targets and TGFB1 mRNA levels across PanCK+ AOIs colored by Subtype class. F. T-SNE plot displaying results of unsupervised clustering of scRNA seq data from eight non-muscle invasive tumors. G. T-SNE plot showing the results of epithelial subset re-clustering. Top panel: Plot on the left is colored by Seurat clusters and plot on the right is colored by sample ID. Bottom panel: Cells colored by luminal (left) and basal (right) gene marker expression. H. Left: T-SNE plot showing the results of epithelial subset re-clustering colored by Subtype assignments. Right: Stacked bar plot indicating the frequency of assigned Subtypes for each sample. I. Dot plot highlighting the pathways significantly enriched in cells within each Subtype relative to the remaining epithelial cells. J. Left: Violin boxplots showing the average expression of MYC, PPARG, IRF4 regulon genes in the cells assigned to each Subtype class. Right: Density plot for CIS Score across the Epithelial cell population.

Subtype3 had enrichment for transcription factors involved in DNA damage response and cell cycle regulation, including BRCA1, CDKN2A, and CENPA. Subtype4 tumors displayed significantly higher MYC regulon activity, consistent with MYC’s role as a master ribosome biogenesis and protein synthesis regulator^29,30^. These findings underscore the interpatient heterogeneity driven by distinct transcriptional programs bladder cancer and support regulon analysis to infer biological function. We independently validated the biology identified by our classification in a previously reported cohort of non-muscle invasive bladder tumors (Zuiverloon 2023 cohort)^31^. We find hallmarks of each Subtype to be recapitulated in this independent cohort, with PPARG downregulation observed in Subtype2 tumors, elevated CIS score in Subtype3 tumors, and higher FGFR3 mRNA levels and MYC regulon activity in Subtype4 tumors **(Supp Fig 1D**).

### Multi-omics transcriptional validation of subtypes by spatial and single-cell profiling to investigate heterogeneity

A potential limitation of bulk RNA sequencing is the potential averaging of expression profiles from individual cells across multiple cell types and tissue compartments, resulting in limited resolution of tumor heterogeneity of especially rare but critical cells. Since the proposed subtyping strategy was dependent on cell-type specific targets, we were interested in resolving the heterogeneity of high-risk bladder cancers. Thus, we validated the expression-based subtyping method in two orthogonal sample types that include similar stage (T1) tumors with spatial analysis and single-cell RNA-Seq. To our knowledge, this is the first cross-omics evaluation of bulk RNA-seq-derived gene signatures in bladder cancer. We performed Digital Spatial Profiling (DSP) using the NanoString GeoMx human whole transcriptome atlas^32^ to differentiate tumor-specific expression profiles (epithelial) from the tumor microenvironment (stroma). Using the Subtype gene signatures, we analyzed 129 pan-cytokeratin-positive tumor AOIs from 43 treatment-naive T1 tumors (Fig 2A). Each PanCK+ Area of Interest (AOI) was meticulously selected to reduce stromal contamination, effectively precluding the examination of tumor AOIs with immune infiltration, such as those that might classify as Subtype2. For each patient, we collected a median of 3 AOIs (range: 1-5), to investigate spatial heterogeneity in subtype expression, we limited the analysis to patients for which more than 1 AOIs were profiled. 51% of patients (18/35) did not exhibit heterogeneity in Subtype expression; of these, 44% classified as Subtype4, 28% as Subtype3, and 28% as Subtype1 (**Fig 2A**). The remaining 17/35 patients had a heterogeneous subtype expression across the tumor AOIs (**Fig 2A**). Comparing the gene expression programs between the AOIs in each class, we find the biological processes identified from bulk RNA sequencing analysis were preserved within the tumor AOIs from each Subtype (**Fig 2B**). For example, Subtype1 showed elevated fatty acid metabolism programs, Subtype3 tumors showed high expression of cell cycle markers (E2F targets and G2M checkpoint), and CIS scores, and Subtype4 tumors showed a marked increase in hypoxia pathway and MYC targets (**Fig 2D, 2E)**. We analyzed the PanCK-stromal AOIs captured alongside the tumor AOIs to investigate the crosstalk between epithelium and stroma. With Subtype4 epithelium expressing elevated levels of FGFR3 mRNA within the PanCK+ AOI, the neighboring stroma expressed increased FGF ligands (FGF17 and FGF2)(**Fig 2C**).

We further validated our subtyping at the single-cell level by analysis of single-cell RNA sequencing from eight high-grade tumors. A large fraction of cells in each sample were epithelial (**Fig 2F**). We extracted and re-clustered the epithelial population, identifying 11 unique clusters that expressed luminal markers (**Fig 2G**). Interestingly, a subset of cells in epithelial sub-cluster 1 expressed high levels of luminal and basal markers, a phenomenon that while rare, has been documented previously^6^ (**Fig 2G**). Next, we assigned each cell an identity based on the Subtype gene signatures. Six of the eight tumors analyzed had greater than 50% of cells assigned as Subtype4 with one tumor sample classified as Subtype3. One tumor sample had a heterogenous profile with 40% of cells belonging to Subtype4 and 45% belonging to Subtype1. A small fraction of cells in two samples classified as Subtype2 (**Fig 2H**). Next, the hallmarks of each Subtype identified previously were validated on the single cells that were classified within each group. Epithelial cells classified as Subtype4 had higher expression of TGFB signaling pathway and MYC targets, and Subtype3 cells had a higher expression of DNA repair genes, cell cycle pathway, and IL2 STAT5 signaling. The small subset of cells that were classified as Subtype2 overexpressed markers of EMT and allograft rejection and cells with Subtype1 identity had elevated expression of cholesterol homeostasis and protein secretion pathways (**Fig 2I**). Next, we validated the regulon profiles observed in bulk RNA sequencing data in the Epithelial population in the single cell data and found MYC regulon activity to be significantly higher in Subtype4, PPARG regulon activity to be significantly lower, and IRF4 activity to be higher in cells with Subtype2 identity. Lastly, we find the CIS score to be the highest in the one tumor sample with the largest fraction of cells belonging to the Subtype3 subtype (scRNA16) (**Fig 2J**). In summary, each subtype’s unique gene expression programs and pathways identified using bulk RNA sequencing were validated through isolated interrogation of the epithelium by Digital Spatial Profiling and single-cell RNA sequencing, highlighting heterogenous gene expression profiles of bladder cancer that appear to be retained across cell compartments and at single-cell resolution.

### Genomic diversity in High-Risk Bladder Cancer

To complement the transcriptional profiling, we expanded our analysis by investigating of the somatic mutational profiles of 92 tumors. Consistent with previous reports, we identified mutations in at least one chromatin modifier gene in 93% of tumors^6^, and these mutation frequencies did not differ significantly across subtype (**Fig 3A**). We then profiled the mutations for each Subtype. Subtype1 tumors had the highest frequencies of mutations in ERBB2 (45%) and ERBB3 (48%) (Fisher’s exact test p, ERBB2 = 2.1e-04, ERBB3 = 7.4e-03) (**Fig 3B**). ERBB2 and ERBB3 are members of the EGFR family of tyrosine kinases that activate downstream pathways such as RAS-ERK and PI3K-AKT, playing essential roles in cell proliferation, differentiation, and migration. Subtype2 tumors were characterized by mutations in genes implicated in immune suppression. PPARG mutations were detected in 31% of Subtype2 tumors (Fisher’s exact test p = 4.5e-02), along with XPO1 mutations in 38% of tumors (Fisher’s exact test p = 1.2e-04) (**Fig 3B**). PPARG mRNA was significantly downregulated in Subtype2 tumors relative to other clusters (Kruskal-Wallis p-value = 6.7e-04) **(Supp Fig 2A**). We found that Subtype3 tumors had a higher frequency of mutations in RB1 (55%) (Fisher’s exact test p = 5.6e-03) and TP53 (64%) (Fisher’s exact test p = 7.4e-02) (**Fig 3B**). Consistent with previous reports of TP53 mutations describing a higher frequency in carcinoma in situ (CIS) tumors, we found tumors within Subtype3 to have elevated CIS scores relative to other clusters^33^. **(Supp Fig 2B**).

**Figure 3:**
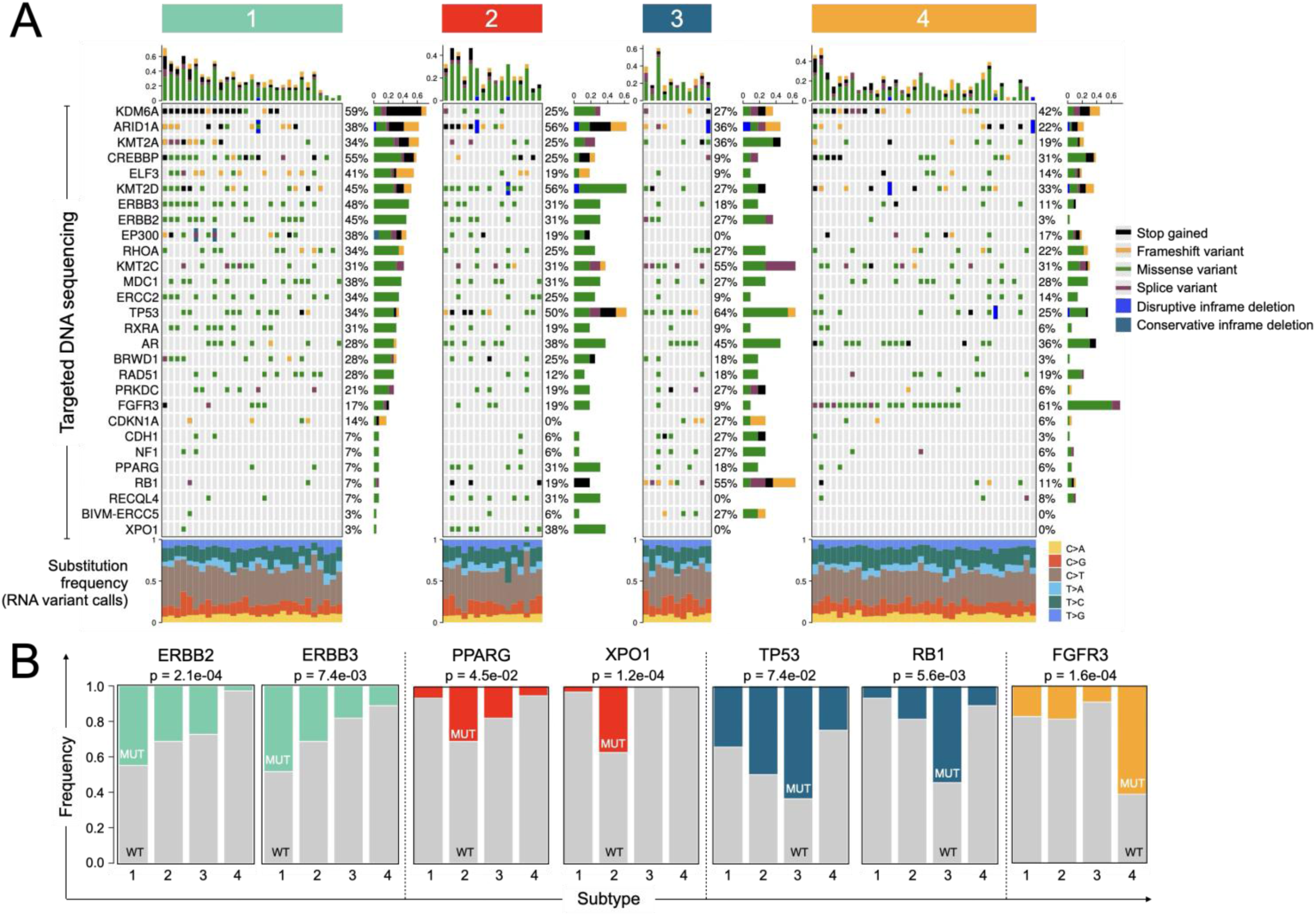
Genomic profiles of unique molecular classes. A. Oncoprint depicting the recurrent somatic mutations identified in tumors within each Subtype (Top). Relative proportions of six possible base-pair substitutions identified by RNA variant calling (Bottom). B. Frequency of specific gene mutations across the Subtypes. (p-values were calculated using a Fisher’s Exact test)

FGFR3 mutations were significantly enriched in Subtype4 tumors (61%) compared to other groups (Fisher’s exact test p = 1.6e-04) (**Fig 3B**). Activating mutations in FGFR3 have previously been associated with the luminal papillary subtype **(Supp Fig 2C**). Consistent with previous findings, we observed a mutual exclusivity pattern between TP53 and FGFR3 mutations^34^ (Data not shown). A feedback loop between FGFR3 and MYC that supports the oncogenic dependency of FGFR3-mutated cell lines has been described in bladder cancer^35^, and we found Subtype4 tumors also exhibited high activity of the MYC regulon and significant enrichment of metabolic programs (**Figure 2A**).

To understand the transcriptomic features driven by significant mutations identified in our cohort, we evaluated the network of differentially regulated gene programs agnostic to Subtype assignment. Activating mutations in FGFR3 were associated with upregulated gene programs related to RNA metabolism and protein synthesis, like those upregulated in Subtype4 tumors **(Supp Fig 2A**). RB1-mutated tumors had upregulated cell cycle markers, consistent with the well-established role of RB1 in cell cycle regulation. TP53-mutated tumors had elevated DNA damage response programs, mitotic regulators, and increased expression of IFN-dependent antiviral genes. XPO1-mutated tumors showed elevated TLR and Interleukin signaling pathways, adaptive immune response genes, genes involved in Rho-GTPase signaling, and a significantly higher inferred immune cell infiltrate based on the ESTIMATE^25^ algorithm **(Supp Fig 2D**).

### RNA editing activity in NMIBC

Mutation analysis of RNA sequencing variant calls can be used to decipher RNA editing processes in tumors that might contribute to tumor heterogeneity^6,36^. To identify mutational processes in high-risk bladder cancers, we employed stringent criteria to identify SNVs from RNA sequencing data and generated single base substitution (SBS) signatures to deconvolute the underlying molecular processes^6,36^. Signatures associated with aging, SBS1, and SBS5, were detected in most tumors (98% and 99%, respectively) (**Fig 4A**). Yet, other signatures were enriched in specific Subtypes. SBS84, proposed to be caused by activation-induced cytidine deaminase (AID)^37^ activity associated with B-cell rearrangement, was found in 43% of Subtype2 tumors (10/23) and was significantly higher in tumors with a high ESTIMATE-derived ImmuneScore (**Fig 4B**).

**Figure 4:**
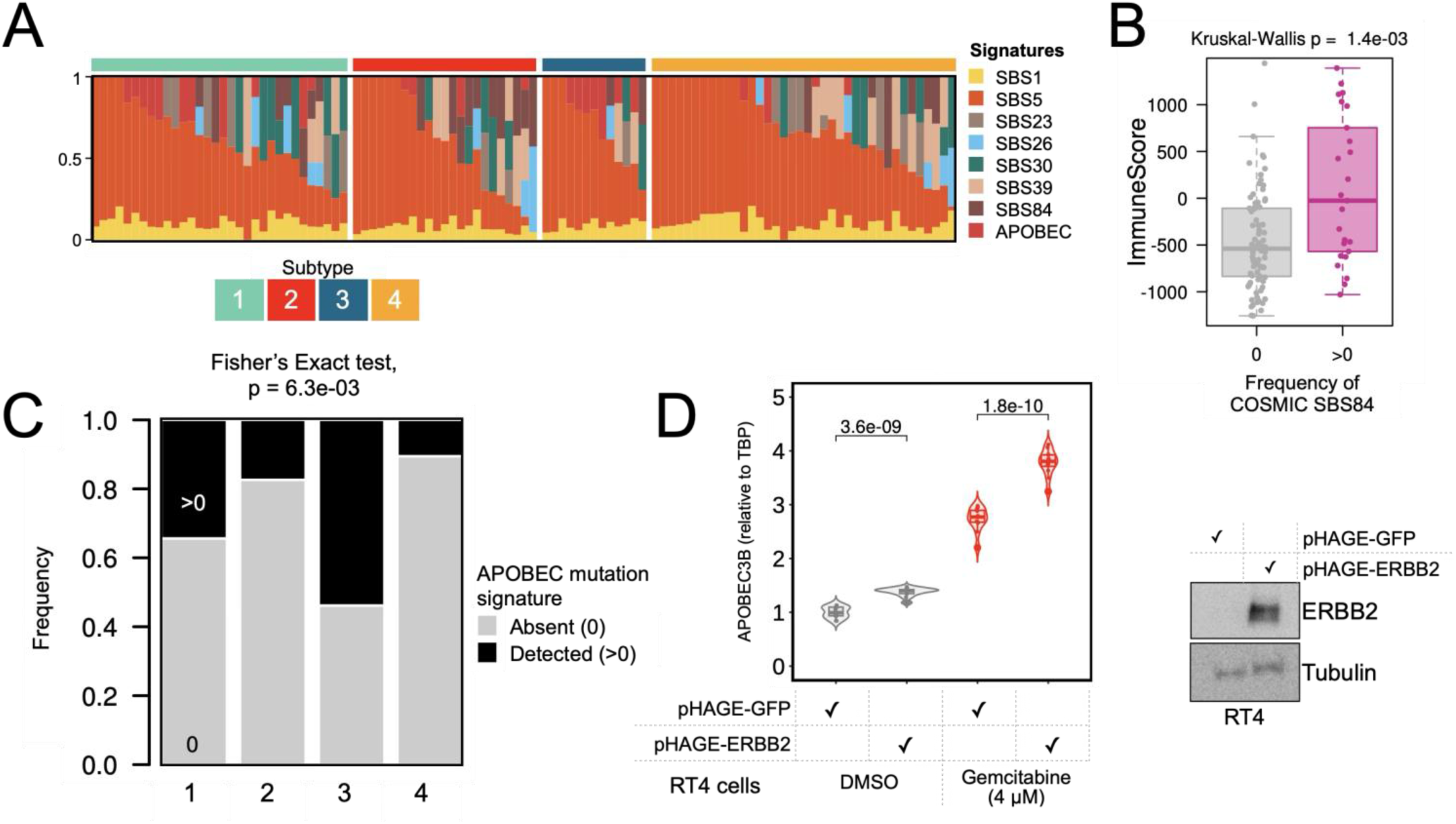
RNA editing activity in high-risk non-muscle invasive bladder cancer. A. Relative contributions of RNA derived COSMIC signatures by Subtypes. SBS2 and SBS13 were combined to define one signature, labeled as APOBEC. B. Distribution of Estimate-derived ImmuneScore by presence/absence of COSMIC SBS84 signature. (p-value was calculated from a two-sided Kruskal-Wallis test) C. Relative contributions of COSMIC APOBEC signature (SBS2, SBS13) by Subtypes. (p-value was calculated using a Fisher’s Exact test) D. APOBEC3B mRNA levels in RT4 cells overexpressing GFP or ERBB2, treated with DMSO or 4μM Gemcitabine for 24 hours (left). ERBB2 protein levels in RT4 cells overexpressing control-GFP or ERBB2 (right). (p-values were calculated from a two-sided Student’s t-test.) (n=2)

An estimated 67% of Stage II disease (MIBCs) mutations are associated with APOBEC mutagenesis, described by SBS2 and SBS13 COSMIC signatures^22,38,39^. We combined SBS2 and SBS13 to create a composite APOBEC signature that was detected at a significantly higher frequency in Subtype3 (53%) and Subtype1 (34%) tumors (Fisher’s exact test p = 6.3e-03) (**Fig 4C**). While the increased APOBEC signatures in Subtype3 may be secondary to genomic instability from the enrichment of TP53/RB1 mutations, we hypothesize that APOBEC signatures found in Subtype1 may be secondary to ERBB2/3 mutations. ERBB2 mutations have been associated with APOBEC mutagenesis in breast cancer^40^, however, to date, ERBB2 alterations have not been associated with APOBEC processes in bladder cancer. To evaluate if ERBB2 directly regulates APOBEC expression, we overexpressed ERBB2 in the RT4 bladder cancer cell line. With overexpression of ERBB2, we observed a moderate, yet significant increase in APOBEC3B mRNA levels, which was further amplified under conditions of replication stress, such as those observed during tumor development or induced by chemotherapy/gemcitabine treatment^41,42^ (**Fig 4D**). This suggests that ERBB2 alterations in conditions of replication stress, such as those found during early tumor development found uniquely in Subtype 1, may be associated with increased APOBEC activity, contributing to intratumor heterogeneity in high-risk bladder cancers.

### FGFR3 mutated tumors are sensitive to MEK and MYC inhibition

Subtype4 accounted for the largest proportion of tumors within our cohort (36%) (**Fig 7**) and was enriched for mutations in FGFR3 (61%) (**Fig 3A and 3B**), and a significantly higher expression of FGFR3 relative to other clusters **(Supp Fig 2D**). To investigate potential therapeutic targeting of Subtype4 tumors, we utilized publicly available gene expression data for 1840 cancer cell lines from CCLE (Cancer Cell Line Encyclopedia)^43,44^ and GDSC (Genomics of Drug Sensitivity in Cancer)^45^ data for 499 chemical perturbations. We classified cell lines into two groups based on a Subtype4 gene expression program (GEP) score. Notably, cell lines with a high Subtype4 GEP score demonstrated elevated expression of FGFR3 mRNA (Kruskal-Wallis p = 2.01e-16) (**Fig 5A**). Leveraging the corresponding GDSC data for these cell lines, we identified a subset of drugs that exhibited increased sensitivity (lower IC50) in cell lines with elevated Subtype4 expression. Several tyrosine kinase inhibitors targeting MEK EGFR were among the top hits. EGFR hyperactivity was recently identified as a mechanism of resistance to FGFR inhibition in bladder cancer^46–48^. MEK is one of the downstream effector pathways activated upon FGFR3 ligand binding (**Fig 5B**). To validate this finding in vitro, we overexpressed FGFR3 S249C mutations in two bladder cancer cell lines, UMUC3 and UMUC9. Upregulation of FGFR3 S249C resulted in higher expression of FGFR3 and upregulation of MYC protein (**Fig 5C**). Furthermore, cell lines with overexpression of the FGFR3 S249C mutation displayed increased sensitivity to Trametinib, a MEK inhibitor, and Myci975, a recently described MYC inhibitor^49^ (**Fig 5D**). These findings identify MEK and MYC as promising targets for mono- or combination therapy with FGFR inhibitors in bladder cancer patients with elevated Subtype4 gene expression. Additionally, MEK or MYC inhibitors may be a viable rescue strategy in FGFR3-mutated tumors that develop resistance to FGFR inhibitors, such as TAR-210

**Figure 5:**
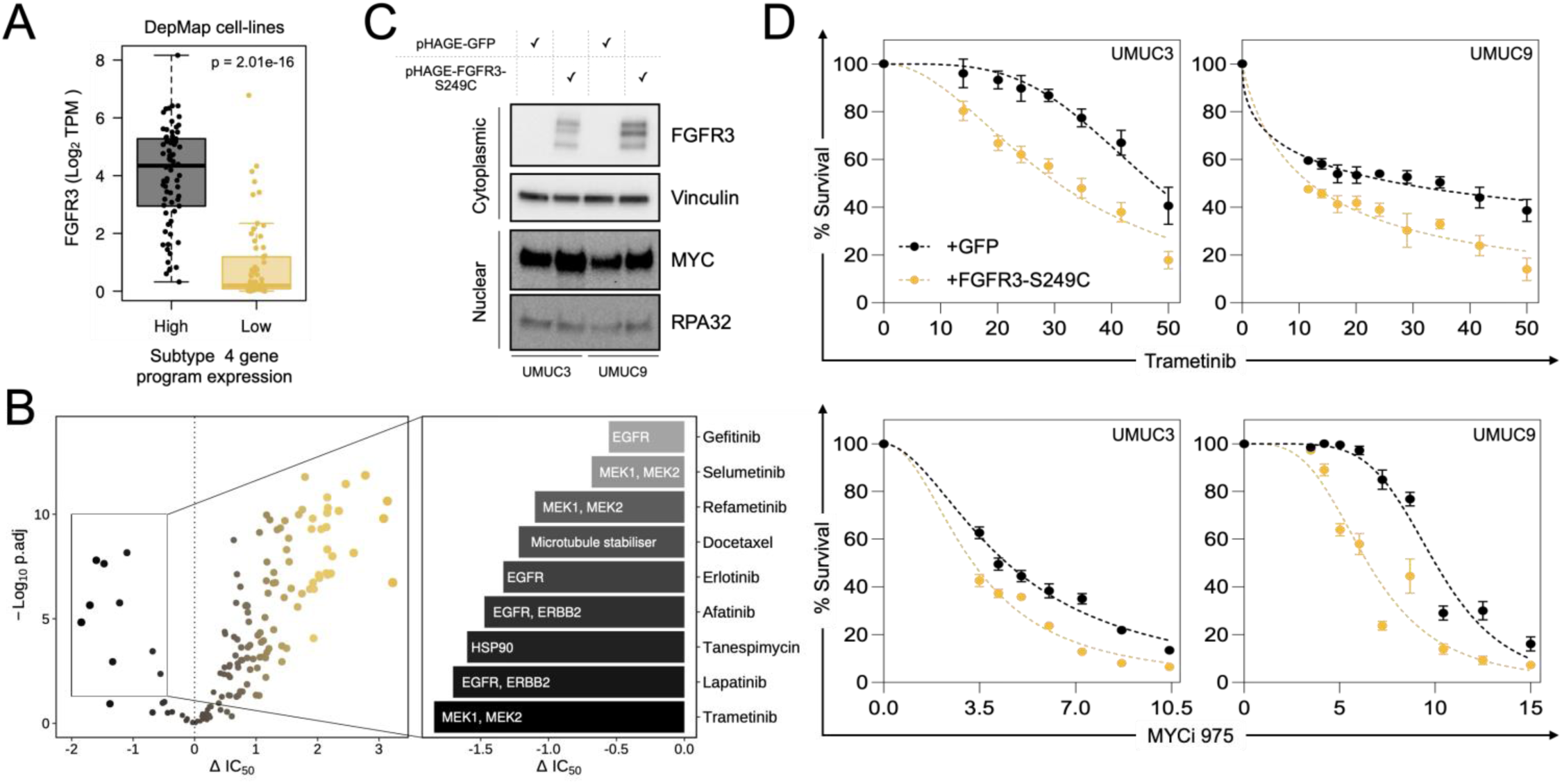
Invitro screen to identify drug targets for Subtype4 tumors. A. FGFR3 mRNA levels across DepMap cell lines classified as having high or low expression of Subtype 4 gene program markers. (p-value was calculated from a two-sided Kruskal-Wallis test) B. Volcano plot showing differences in IC50 between cell-lines with high vs low Subtype 4 gene program score on the x-axis and −Log10 adjusted p-value for each comparison on the y-axis. Each point represents a unique chemical perturbation (Left). Drugs with increased sensitivity in cell-lines that have high expression of Subtype 4 gene program (Right). C. MYC and FGFR3 protein levels in UMUC3 or UMUC9 cell-lines overexpressing GFP controls or FGFR3 S249C mutations. (Vinculin was used as loading control for the cytoplasmic fraction, RPA32 was used as loading control for the nuclear fraction) (n=2) D. Survival curves for UMUC3 and UMUC9 cell-lines overexpressing GFP controls or FGFR3 S249C mutations treated with MEK inhibitor Trametinib or MYC inhibitor for 7 days. (n=3).

### Differentiating characteristics of Subtype2 tumors

Subtype2 tumors had the longest recurrence-free survival to BCG and were found to have enhanced markers of anti-tumor immunity with higher infiltration of B cells, NK cells, CD8+ T cells, macrophages, and myeloid dendritic cells (**Fig 6A**). Since these tumors appeared to have higher baseline levels of immune infiltration before BCG, we were interested in the possible mechanisms of enhanced immunity. These possible features included tumor intrinsic features such as diminished PPARG activity **(Supp Fig 2A**) and tumor microenvironment features that increase the immune response.

**Figure 6:**
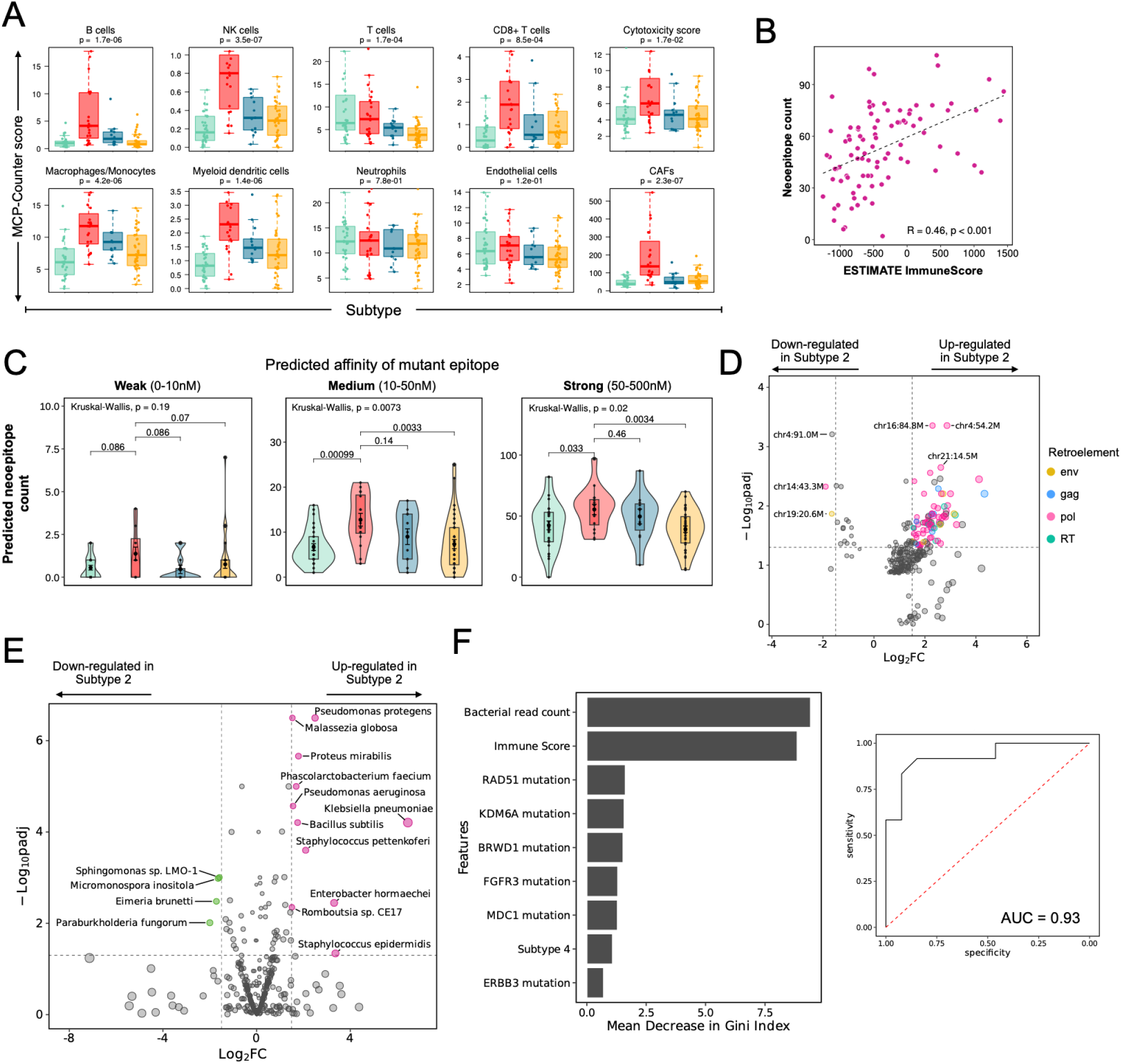
Unique features of immune-infiltrated tumors. A. Boxplot highlighting the population abundances of tumor infiltrating immune and stromal cells estimated by MCP-Counter algorithm across Subtypes. (p-values were calculated from a two-sided Kruskal-Wallis test) B. Correlation between the neoepitope count and the ESTIMATE-predicted immune cell fraction. (R indicates Pearson correlation coefficient) C. Distribution of neoantigen frequency by predicted affinity for MHC across Subtype tumors. (p-value was calculated using a two-sided Kruskal-Wallis test and for individual comparisons using a two-sided Wilcox test) D. Volcano plot comparing expression of endogenous retroviral elements (ERV) in Subtype2 tumors vs the rest of the cohort. E. Volcano plot comparing expression of microbial species in Subtype2 tumors vs the rest of the cohort. F. Left: Bar plot displaying the relative importance of each predictor in the final random forest model. Right: ROC curve depicting the performance of the random forest model.

#### Enhanced antigen presentation

A high neoantigen burden has often been associated with enhanced anti-tumor immunity^50^. To address whether the difference in pre-existing anti-tumor immunity between the Subtypes resulted from differences in neoantigens presented by the tumor, we developed a pipeline integrating HLA allele typing and MHC binding predictions for tumor-specific mutant peptides. While our analysis was limited to the genes targeted within our custom panel, we identified an average of 53 neoepitopes per tumor (Range: 2-107, median: 53.5). We observed a statistically significant positive correlation (Pearson coefficient = 0.46, p < 0.001) between tumor neoepitope count and the immune cell fraction inferred by ESTIMATE (**Fig 6B**). We further categorized the neoantigens into three classes based on the predicted affinity of the neoepitope for MHC: Weak binders (0-10nM), Medium binders (10-50nM) and Strong binders (50-500nM). Notably, Subtype2 tumors had a consistently higher frequency of neoantigens with a strong affinity for MHC (**Fig 6C**).

#### Tumor microenvironment

We characterized the molecular features that could modulate the pre-treatment immune microenvironment. Based on descriptions of enhanced immunity in the skin associated with commensal bacteria and local activation of ERVs^51^, we hypothesized that re-activation of endogenous retro-elements^52^ or exposure of the bladder epithelium to bacteria or viruses before BCG therapy contributes to the recruitment of immune cells to the bladder. To investigate this hypothesis, we compared the expression of transposable elements such as long interspersed nuclear elements (LINE) and endogenous retroviral elements (ERV)^51^ in tumors with higher than median Immune Score (measured by ESTIMATE). We found elevated expression of several endogenous retroelements in Subtype2 tumors (**Fig 6D**). Additionally, we performed meta-transcriptomic deconvolution^53^ and identified significant upregulation of several bacterial species known to cause urinary tract infection within Subtype2 tumors (**Fig 6E, Supp Fig 3A**). Overall, our results indicate that NMIBC tumors with a potent pre-existing anti-tumor immunity might be a good candidate for BCG

### Integrated multi-factorial model of Response to BCG Immunotherapy

Our study identified multiple factors that can influence BCG response in T1 tumors: mutational and transcriptomic profile of a tumor, presence of APOBEC mutation signature, tumor mutation burden, neoantigen burden, density of immune cell infiltrates, and microbial populations. However, relationships between these variables may contribute to the response to BCG, and these remain largely uncharacterized. The ability to predict BCG response holds significant clinical implications, prompting us to explore various machine learning algorithms for developing a predictive model, each presenting unique strengths and tradeoffs. Through extensive testing involving combinations of 28 features, we constructed a random forest model for BCG response prediction (**Fig 6F**). The best model prediction was obtained using a combination of nine features which included density of bacterial read count, immune score, subtype 4 status, and mutations in RAD51, KDM6A, BRWD1, FGFR3, MDC1 and ERBB3 (AUC = 0.93, **Fig 6F**).

## Discussion

BCG is the primary immunotherapy for high-risk bladder cancer for 50 years. Yet, predicting the response to BCG is challenging because the anti-tumor activity is not explicitly targeted towards any genomic characteristics. While over 800,000 vials of BCG are produced annually, the unpredictable distribution shortages of BCG limit its broad use for the foreseeable future. Thus, investigating alternative therapies is an urgent and unmet clinical need. This study investigated the genomic and expression-based features of treatment-naïve high-risk bladder cancer to identify the tumor and tumor microenvironment. We describe four distinctive and potentially targetable meta-clusters by harmonizing transcriptomic analysis with matched targeted sequencing data. (**Fig 7**). Finally, we develop a machine learning-based model to parse these individual features and identify those associated with clinical recurrence.

**Fig 7:**
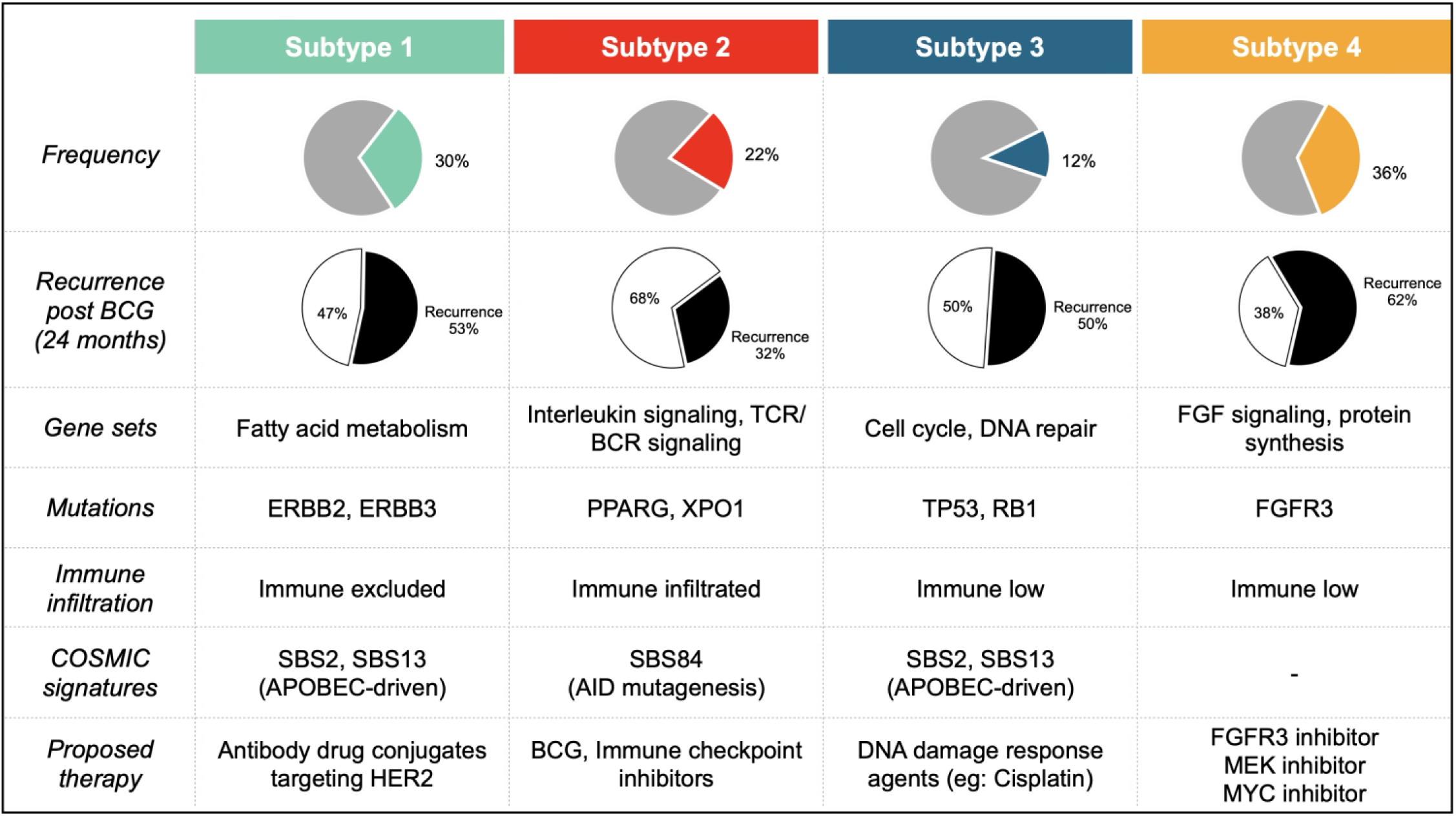
Summary characteristics of Subtype NMIBC subtypes.

The hallmark of Subtype1 was mutations in ERBB2/ERBB3 and a subset of tumors with ongoing APOBEC activity. In a retrospective analysis of 454 NMIBC patients, Tan et al. identified high HER2 expression associated with poor response to BCG^54^. The application of antibody-drug conjugates for targeted bladder delivery offers a potential treatment for this Subtype. The distinctive genomic landscape of Subtype3 was marked by a genomic instability with an increased prevalence of TP53/RB1 mutations, heightened activity of cell cycle and DNA repair pathways, and increased APOBEC mutagenesis. Based on the hallmarks of replication stress, DNA-damaging agents, such as single or multi-agent intravesical chemotherapy such as mitomycin C, gemcitabine/TAR200, or docetaxel may be the most effective. The distinct signature of Subtype4 is enriched FGFR3 mutations, concomitant with increased MYC regulon activity. In addition to approved FGFR3 inhibitors such as erdafitinib or TAR210 device, we identify two other therapies, targeting MYC and MEK, as potential therapies for Subtype4 either as monotherapy or in combination with FGFR3 inhibitors^55^.

Subtype2 tumor had increased immune infiltration, enhanced inflammatory gene sets and cytokines at baseline before BCG. Remarkably, patients within Subtype2 had the most favorable prognosis, with an impressive 86% maintaining a recurrence-free status at 12 months. Additionally, we identified the re-activation of several endogenous retroelements in tumors with a higher immune infiltrate and increased expression of bacteria and viruses. This Subtype appears to be the most responsive to BCG and ensuring that Subtype 2 tumors receive BCG could further improve clinical response.

Prior reports have underscored the association between asymptomatic bacteria and improved response to BCG therapy^56–58^. Patients diagnosed with asymptomatic bacteriuria before receiving BCG treatment experience longer recurrence-free survival^57^. In parallel, administering antibiotics before or during BCG or systemic checkpoint therapy is associated with worse recurrence-free survival^56,59^. These findings highlight the vital need to investigate the bladder tissue-associated microbiota interaction in modulation of the response to immunotherapy. Direct manipulation of the bladder microenvironment is currently under investigation to modulate the immune response.

Our study has several limitations that deserve attention. First, the sample size of our cohort remains limited. Since a significant fraction of our analysis, including but not limited to the final random forest model, requires access to raw sequencing data, we could not validate our predictive model in other publicly available datasets.

Due to the limited access of BCG immunotherapy, new therapeutics are critical to improving survival of patients with high-risk bladder cancer. We identify a new expression-based system to target unique drivers in. high-risk bladder cancer, and identify their essential role in single cells and in spatially in tissue compartments. Our research identifies a specific patient subgroup (Subtype2) that will have the most benefit from BCG therapy and we nominate multiple mechanisms that can regulate the local immune microenvironment. In other subtypes, we identify genetic and expression-based susceptibilities that may be targeted with available therapies not currently used to treat bladder cancer.

## Materials and methods

### Patient cohorts, IRB, and clinical data

Approval from the Institutional Review Board (IRB) was obtained to assess a group of patients with advanced stage T1 bladder cancer at Northwestern Memorial Hospital. To conduct the study, the Northwestern Institutional Review Board (STU00204352) granted permission to waive the requirement for informed consent. All patients included in the study were newly diagnosed with bladder cancer and had not received prior intravesical chemotherapy or immunotherapy treatment. The researchers recorded the dates of transurethral resection (TUR), repeat TUR, and the amount of Bacillus Calmette-Guérin (BCG) administered to each patient. All patients in the study completed at least the initial phase of BCG treatment, which consisted of six doses at full strength. Recurrence was defined as development of a high-grade bladder tumor after the last TUR. Response was defined as recurrence after 24 months of disease free status.

### FFPE RNA extraction, library construction, and sequencing

A GU pathologist annotated tumor sections for each FFPE slide. These sections were macrodissected and processed for RNA and DNA extraction. Genomic DNA and RNA were extracted from macrodissected FFPE tissues using a Mag-Bind FFPE DNA/RNA 96 kit (Omega Biotek, Norcross, GA, US) following the manufacturer’s recommendations. Isolated RNA sample quality was assessed using the High Sensitivity RNA Tapestation (Agilent Technologies Inc., California, USA) and quantified by Qubit 2.0 RNA HS assay (ThermoFisher, Massachusetts, USA). Ribosomal RNA depletion was performed with Ribo-Zero Plus rRNA Removal Kit (Illumina Inc., California, USA). Subsequently, libraries were constructed using Truseq Stranded Total RNA kit (Illumina, California, USA). Final library quantity was assessed by Qubit 2.0 (ThermoFisher, Massachusetts, USA), and quality was assessed by TapeStation D1000 ScreenTape (Agilent Technologies Inc., California, USA). Equimolar pooling of libraries was performed based on QC values and sequenced on an Illumina NovaSeq (Illumina, California, USA) with a read length configuration of 150 PE for 80 M PE reads per sample (40M in each direction).

### Gene expression analysis

RNA seq reads were trimmed using Trimmomatic v0.39^60^ and aligned to the human genome GRCH38 v103 using STAR v2.7.9a^61^ for both cohorts. Next HTSeq v2.0.1^62^ was used to compute raw gene expression counts from aligned reads. Additionally, Salmon v1.5.0^63^ was used to quantify transcript level expression data to use as an input for immune deconvolution analysis. Next, to allow for processing of data generated in both cohorts, we performed batch correction using the counts file generated using HTSeq counts file as an input to Combat-seq function within sva package v3.42.0^64^. To generate FPKM-normalized gene expression data, we used batch corrected raw counts as an input to DESeq2 v1.34.0^65^.

### Clustering analysis

To identify the ideal number of clusters using transcriptomic data, we performed consensus clustering using ConsensusClusterPlus v1.58.0^66^, with the top 1500 genes ranked based on median absolute deviation (MAD) as an input, using pearson correlation for the distance metric and hierarchical clustering algorithm with 10,000 resampling repetitions. We determined the optimal k=4 and computed average silhouette using R package CancerSubtypes v1.20.0^67^. To classify tumors within our cohort using previously described approaches, we used FPKM expression data as an input to classifyT1BC v0.2.0^20^, BLCAsubtyping v2.1.1^68^, classifyNMIBC v1.1.0^6^ and consensusMIBC v1.1.0^68^. Classification of tumors in our cohort using each subtyping method was visualized using a Sankey plot generated using networkD3 v0.4. All heatmaps were generated using ComplexHeatmap v2.10.0^69^.

### Differential gene expression analysis and Immune deconvolution

Differential gene expression analysis was conducted using DESeq2 v1.34.0 using batch corrected raw counts as an input. Next genes were filtered based on FDR<0.05 and .rnk files containing significant genes and the Wald statistic values were generated using the differential expression data and used as an input to GSEA (Gene Set Enrichment Analysis) app v 4.3.2 for pathway analysis^70^. Pathways used were defined by the geneset file Human_AllPathways_January_01_2023_symbol.gmt downloaded from the Bader Lab website. The pathway analysis results were visualized using EnrichmentMap v3.3.6^71^ within the Cytoscape app v3.10.0^72^ which connects pathways with shared gene names using a Markov Cluster algorithm. FPKM expression data was used as an input for the immunedeconv R package v2.1.0^73^, to evaluate immune cell infiltration based on RNA expression data using two different algorithms: ESTIMATE and MCPCounter^25,73–75^.

### Regulon analysis

We used RTN package v2.18.0^26,76^ to generate a regulatory network using 2708 transcription factors. We applied 250 bootstraps to generate the transcriptional network. Next using the tni.gsea2 function within the RTN package, we calculated regulon activity profiles for regulons with a minimum size of 15 “posORneg” targets. Next, we performed transcriptional network analysis for each Subtype using the rtni object generated in the previous step and a vector of differentially expressed genes. Networks for each Subtype were filtered based on log2FoldChange > 1.2 and padj < 0.05 and the resulting regulon gene list was used to generate a binary tree-and-leaf diagram that combines tree and force-directed layout algorithms using the TreeAndLeaf package v1.6.1^77^. Boxplots were generated using boxplot() function in R.

### DNA library construction and sequencing

All DNA isolates were quantified using Qubit 2.0 DNA HS Assay (ThermoFisher, Massachusetts, USA). Pre-hybridization library was produced using SureSelect XT HS and XT low input library preparation kit (Agilent Technologies, CA, USA) and barcoding was performed using Illumina 8-nt dual-indices. The hybridization was performed with customized IDT discovery pool based on manufacture’s protocol. The enriched libraries were quantified using Qubit 2.0 DNA HS Assay and library quality was evaluated by TapeStation HSD1000 ScreenTape (Agilent Technologies, CA, USA). Enriched libraries were pooled before sequencing on an Illumina NovaSeq S4 sequencer for 150 bp read length in paired-end mode.

### DNA data analysis

Adapter sequences and low-quality bases were trimmed from Fastq files using trim_galore v0.6.4_dev and FastQC v0.11.9 was performed on the trimmed reads. The trimmed reads were mapped to human reference genome version hg38 using bwa-mem v0.7.17. Next, duplicate reads were marked using markduplicate utility from Picard (GATK4) v4.1.9.0 tool, where duplicate reads are defined as originating from a single fragment of DNA. Coverage profiles were generated using mosdepth v0.3.1. Finally, analysis ready BAM files were generated by excluding reads with MAPQ < 30 with samtools v1.11^78^. Variant detection was performed using Pisces tool and resulting variants were annotated using SnpEff and SnpSift v5.0^79,80^. Annotated variants were filtered to retain high-quality somatic variants using several criteria including but not limited to variant allele frequency, functional annotation, population allele frequency annotation and variant quality. VCF files containing annotated variants were imported into R using vcfR package v1.13.0. Oncoplots were generated using genes mutated in at least 10% of the cohort using the oncoPrint function in ComplexHeatmap package v2.10.0^69^.

### RNA-based variant calling and mutation signature analysis

We used the GATK pipeline^81–83^ (gatk v4.1.0) for identifying single nucleotide variants in RNA seq data. The process involved aligning the raw RNA reads to the GRCH38 v103 human genome assembly using STAR v2.7.9a^61^ using two-pass mode, marking duplicates with PICARD tools v 2.27.4, followed by further processing using SplitNCigarReads, BaseRecalibrator, and ApplyBQSR to reformat alignments spanning introns and correct base quality scores. Variants were then called using the HaplotypeCaller software and converted to vcf files using GenotypeGVCF. The resulting VCF files were annotated using SnpEff v4.3t^79^. Variants were filtered using SnpSift^80^ to keep variants with a quality score above 100, a Fisher Strand score (FS) below 30.0, and a HIGH or MODERATE impact. VCF files containing filtered variants for each sample were used to identify mutational signatures using SigProfilerExtractorR v1.1.13^84^ with the following criteria: GRCH38 reference genome, normalization method = “gmm”, nmf_replicates = 500, cosmic_version = 3.3. For each sample, fraction of contribution of each COSMIC signature was identified. Mutation count identified for SBS2 and SBS13 were summed to generate a composite APOBEC signature.

### Drug sensitivity analysis

To identify drugs that might show unique sensitivity to cell-lines with elevated expression of a particular Subtype program, we downloaded drug response data for 499 compounds publicly available from GDSC (Genomics of Drug sensitivity in cancer) website^45^. Drug sensitivity data was matched to TPM expression data available for cell-lines within the Cancer Cell Line Encyclopedia (CCLE) from the Dependency Map (DepMap) portal^43,44^. After filtering, IC50 data for 499 compounds across 695 cell-lines was used for the final analysis. A Subtype4 program score was generated for each cell-line, next cell-lines were ranked based on the program score and cell-lines in the top 10% and bottom 10% quantile were compared for IC50 analysis.

### Invitro experiments and drug treatment

UMUC3 or UMUC9 cell-lines were used for all in-vitro experiments described. pHAGE-GFP was a gift from Jay Shendure (Addgene plasmid# 106281)^85^, pHAGE-FGFR3-S249C (Addgene plasmid# 116386)^86^ and pHAGE-ERBB2 (Addgene plasmid# 116734)^86^ were a gift from Gordon Mills & Kenneth Scott. Plasmids were packaged into lentiviral particles using lentiviral packaging plasmids psPAX2 and pm2D.G, both plasmids were a gift from Didier Trono. (Addgene plasmid# 12260 and 12259). Stable cell-lines were generated and selected using flow cytometry for expression of GFP marker or puromycin. Cell-lines were treated with Trametinib or Myc inhibitor (Myci975^49^) at indicated concentrations for seven days post which survival was assayed for survival using Resazurin absorbance as a readout. For experiments involving pHAGE-GFP and pHAGE-ERBB2 overexpression, cells were treated with DMSO or Gemcitabine (4µM) for 48 hours, post which cells were harvested for RNA extraction. RNA was extracted using RNeasy Mini Kit (Qiagen 74104), and cDNA was generated using High-capacity cDNA reverse transcription kit (ThermoFisher Scientific, 4368814). Quantitative PCR was performed using Luna Universal qPCR master mix (NEB, M3003) and a BioRad CFX Connect Real Time System. Protein expression was verified using western blotting with these antibodies: anti-ERBB2 (Cell Signaling Technologies, 2165), anti-FGFR3 (Cell Signaling Technologies, 4574), anti-RPA32 (Cell Signaling Technologies, 35869), anti-Tubulin (Sigma, T5168), anti-Vinculin (Invitrogen, MA5-11690), anti-MYC (Invitrogen, 13-2500).

### Retroelement analysis

RNA seq reads were trimmed using Trimmomatic v0.39^60^ and aligned to the human genome GRCH38 v103 containing retroelement nucleotide sequences downloaded from http://geve.med.u-tokai.ac.jp/ using STAR v2.7.9a for both cohorts. Next HTSeq v2.0.1 was used to compute raw gene expression counts from aligned reads using the annotation .gtf file provided in the gEVE database^87^. Next, to allow for processing of data generated in both cohorts, we performed batch correction using the counts file generated using HTSeq counts file as an input to Combat-seq function within sva package v3.42.0. To generate FPKM-normalized gene expression data, we used batch corrected raw counts as an input to DESeq2 v1.34.0^65^.

### Microbial taxonomy classification using KRAKENUniq

Microbiome analysis was performed using KRAKEN software suite. Briefly, host sequences that do not align to GRCh38 v103 and CHM13 human genome sequence were identified using STAR. Next, Non-human reads were used as an input to KRAKENUniq and mapped against a custom database containing bacterial, viral, archaeal genomes downloaded from https://benlangmead.github.io/aws-indexes/k2. Classification report obtained as an output from KrakenUniq was used as an input to Pavian Shiny App v1.2.0 to generate a text file with raw reads from distinct species and genera, which was used as an input for further analysis. Percentage of reads belonging to the microbial species were calculated as a fraction of total sequencing read depth of each sample.

### Immunohistochemical analysis

Formalin fixed paraffin embedded (FFPE) slides from bladder tumors were used for IHC staining for CD45 and LPS. Deparaffinization was done by placing the sections in xylene for 15 minutes, then rehydration through a graded series of ethanol (100%, 95%, 80%, and 70%) and rinsing with distilled water. Antigen retrieval was performed by treating the sections with a citrate buffer (pH 6.0) and heating in a pressure cooker for 10 minutes. Endogenous peroxidase activity was quenched by incubating tumor sections with 3% hydrogen peroxide for 10 minutes. Next, the sections were blocked with 5% bovine serum albumin to minimize nonspecific binding. Primary antibodies targeting the protein of interest were applied overnight at 4°C. After washing with TBS-T, the sections were incubated with a secondary antibody conjugated with HRP for 1 hour at room temperature. Visualization was achieved using 3,3’-diaminobenzidine (DAB) as the chromogen, and the sections were counterstained with hematoxylin. Finally, the slides were dehydrated, cleared, and mounted using a mounting medium. Primary antibodies used in this experiment were anti-CD45 (Abcam ab10558) and anti-LPS (HycultBiotech #HM6011)^88^.

### Digital Spatial Profiling

Digital Spatial Profiling (DSP) procedures were conducted following established methodologies^32^. Slides were deparaffinized by incubating in a drying oven at 60°C for two hours, followed by sequential washes in xylene (3 x 5 min), 100% ethanol (2 x 5 min), 95% ethanol (2 x 5 min), and 1x PBS (1 x 1 min). Target retrieval involved placing slides into a steamer with DEPC-water heated to 99°C for 10 seconds, followed by 1x Tris-EDTA for 20 minutes. Slides were then washed in 1x PBS (5 min) and incubated in 1µg/mL Proteinase K (Thermo Fisher, Waltham, MA) at 37°C for 20 minutes. Post-fixation slides were incubated in 10% neutral buffered formalin (NBF) for five minutes, followed by NBF stop buffer (0.1M tris, 0.1M glycine in DEPC-treated water, 2 x 5 min), and 1x PBS (1 x 5 min).

RNA probe library in situ hybridization was performed using the NanoString GeoMX™ Whole Transcriptome Atlas (WTA) (NanoString, Seattle WA), diluted per manufacturer instructions. Hybridization solution was added to each slide, covered with Grace Bio-Labs Hybrislip™ (Bend, OR) hybridization covers, and incubated in a hybridization oven at 37°C for 20 hours. Stringent washes (100% formamide in equal parts 4X-SSC) at 37°C (2 x 25 min) and 2X SSC wash at room temperature (2 x 2 min) were performed to remove off-target probes. Slides were then blocked using blocking buffer W (NanoString, Seattle, WA) for 30 minutes in a humidity chamber at room temperature. To distinguish tumor and tumor microenvironment (TME), slides were stained with immunofluorescent antibodies from the NanoString solid tumor TME morphology kit [Pan-CK for epithelial cells, CD45 for immune cells, and SYTO 83 for nuclear staining] (NanoString, Seattle, WA) for one hour.

DSP was conducted on prepared slides using the GeoMx digital spatial profiler (NanoString, Seattle, WA). After loading and scanning the slides onto the instrument, regions of interest (ROIs) were manually selected for transcriptomic profiling. Each ROI was divided into tumor and stroma segments based on the presence of immunohistochemical morphologic markers (PanCK+ staining for tumor and PanCK-for stroma). DNA oligonucleotides were aspirated from each segment separately and stored in individual wells in a 96-well plate. As per the default NanoString GeoMx™ protocol, Illumina i5 and i7 dual indexing primers were added to the oligonucleotide tags during PCR for unique indexing. PCR purification utilized AMPure XP beads (Beckman Coulter, Indianapolis, IN). Library concentration was measured using a Qubit fluorometer (Thermo Fisher, Waltham, MA), and quality was assessed using a Bioanalyzer (Agilent Technologies Inc., Santa Clara, CA). Sequencing was performed on an Illumina NovaSeqX (Illumina Inc., San Diego, CA).

Following sequencing, fastq files were converted to DCC files using the GeoMx NGS Pipeline v2.3.3.10, and loaded onto the GeoMx instrument, converted into target counts for each segment. Raw counts were filtered based on two criteria: removal of targets detected below the limit of quantitation (LOQ) and elimination of segments with fewer than 50 nuclei. Filtered raw counts were Q3 normalized for comparison across all segments and were employed for subsequent analyses.

PanCK+ AOIs for each patient were averaged to create a pseudo-bulk PanCK (tumor) expression profile, which was then used as to classify tumors using Subtype gene signatures. Differential expression between PanCK+ segments within each Subtype was performed using limma v3.54.2^89^. To characterize the PanCK-Stromal interactions within tumors in each Subtype, we utilized the publicly available CellPhoneDB database. Interactions were filtered based on gene expression within the DSP cohort and based on differential expression between the PanCK+ and PanCK-segments. Chord diagrams depicting the significant tumor-stromal interactions for each Subtype were plotted using circlize v0.4.15^90^.

### Single cell RNA sequencing of NMIBC tumors

Samples were processed as described before. Briefly, eight fresh bladder tumors were collected under IRB STU00204352 from patients undergoing TURBT. Tissue specimens were cut into small pieces and enzymatically digested to produce a suspension of individual cells. Cells were then suspended in a solution containing 0.04% BSA in PBS and loaded onto a 10X Genomics Chromium platform for Gel Bead-In Emulsion (GEM) generation and barcoding using the Chromium Single GEM Single Cell 3’ reagent kit v3.1. Next, libraries were prepared in accordance with the manufacturer’s instructions. The resulting libraries were combined and sequenced on a single lane of an Illumina Novaseq, generating 150-bp paired-end reads. Demultiplexed read sequences were generated in FASTQ format and subsequently processed using the Cell Ranger v4.0.0 pipeline provided by 10X Genomics. Alignment of reads was performed against the GRCh38 genome, and gene counts from GENCODE v32/Ensembl 98 were quantified using Cell Ranger’s counts command to produce feature-barcode matrices. Samples were further analyzed in R using Seurat v5.0.1. Cell identity was assigned using SingleR v2.0.0. Cells with epithelial identity were extracted and underwent normalization and scaling to standardize gene expression profiles using built in functions in the Seurat package. Variable features relevant to the epithelial compartment were identified using a split-apply combine approach. PCA was then performed using the variable features as input followed by construction of nearest-neighbor graphs to capture local neighborhood relationships between cells using the Seurat package’s functions. Using the computed neighborhood relationships, epithelial cells were clustered into distinct groups with a resolution parameter of 0.1. Expression of basal, luminal markers and CIS genes were investigated^22^. Basal markers included CD44, CDH3, KRT1, KRT14, KRT16, KRT5, KRT6A, KRT6B, and KRT6C, while luminal markers comprised CYP2J2, ERBB2, ERBB3, FGFR3, FOXA1, GATA3, GPX2, KRT18, KRT19, KRT20, KRT7, KRT8, PPARG, XBP1, UPK1A, and UPK2. CIS score was calculated using a ratio of previously described CIS.up and CIS.down genes^6^. Each cell was assigned a Subtype identity based on the expression of Subtype signatures. MYC, PPARG and IRF4 Regulon profiles were similarly assayed using genes enriched within each regulon profile in the bulk RNA sequencing data.

### Neoantigen Prediction

Paired end FASTQ files were processed using Trimmomatic v0.39^60^ to remove low-quality bases and adapters. Next trimmed reads were aligned against the GRCh38 human genome using BWA-MEM^91^, and the resulting BAM files were processed using Picard’s MarkDuplicates tool. Variants were identified using GATK Mutect2^81,82^, with a specified .bed file containing the genomic coordinates of the targeted genes. Next, vcf files were annotated using the Variant Effect Predictor tool (VEP)^92^ from Ensembl. Expression data was added to the annotated VCF files using vcf-expression-annotator function within vatools package. Resulting annotated VCF files were then used as an input to pvacseq function within the antigen prediction toolkit pVACtools v4.0.6^93,94^. Trimmed RNAseq fastq files were used as an input to arcasHLA^95^ to determine the HLA genotype for each sample which was then used as an input to pvacseq. Mutant peptides of different amino acid lengths were generated for the predicted MHC class I (8, 9, 10) and MHC class II alleles (14,15,16,17) for each identified gene mutation. Strength of the MHC:peptide binding affinity for each neoepitope was predicted using NNalign^96^, NetMHC, NetMHCIIpan^97^, NetMHCcons, NetMHCpan, PickPocket^98^, SMM, SMMPMBEC, SMMalign^99^. Median values across all prediction algorithms were used as a composite score of the MHC:peptide binding affinity for each predicted neo-epitope. Next, all predicted neoantigens were classified based on binding affinity^100^ (IC50) into three groups: Weak binders (0-10nM), medium binders (10-50nM) and strong binders (50-500nM).

### Random forest model

The dataset was partitioned into a discovery set (n = 68) and a validation set (n = 21). Eighteen variables, including ERBB2, ERBB3, FGFR3, PPARG, TP53, RB1, XPO1 mutation status, PPARG and MYC regulon activity, APOBEC mutation burden, neoantigen burden, percent of non-human reads, mean expression of ERV (endogenous retroviruses), and assignment to Subtype1, Subtype2, Subtype3, Subtype4, were selected as predictors. The model for response prediction at 24 months was developed using the R package caret (v6.0.95)^101^. To identify the optimal combination of predictors, we employed a stepwise approach. For each selected combination, a random forest model was trained using the train function from the caret package with repeated cross-validation (method = ‘repeatedcv’, number = 100, repeats = 3). The area under the receiver operating characteristic curve (AUC) and accuracy metrics were calculated for each model on the validation set. Additionally, the p-value for comparing each model’s confusion matrix to the null hypothesis of random classification was obtained using the chi-square test. The DeLong method was used to generate p-values for comparing the univariate model to the final random forest model.

## Author contributions

**Conceptualization:** J.J.M.

**Methodology:** K.M., N.F., Y.Y.

**Software:** K.M., J.J.M.

**Validation:** K.M., Y.Y., N.F.

**Formal Analysis:** J.J.M., K.M., N.F., E.Z.L., Y.Y.

**Investigation:** B.C.,J.J.M., K.M., N.F., E.Z.L., Y.Y.

**Resources:** J.J.M., K.M.

**Data curation:** J.J.M., K.M., N.F., E.Z.L., Y.Y.

**Writing-original draft:** J.J.M., K.M.

**Writing-review and editing:** J.J.M., K.M., N.F., S.A.A., B.C., Y.Y., E.Z.L.

**Visualization:** K.M., Y.Y.

**Supervision:** J.J.M., S.A.A.

**Project administration:** J.J.M.

**Funding acquisition:** J.J.M.

