## Supplemental files for "Genomic and transcriptomic profiling of high-risk bladder cancer reveals diverse molecular and microenvironment ecosystems"

**Supplementary Figure 1:**

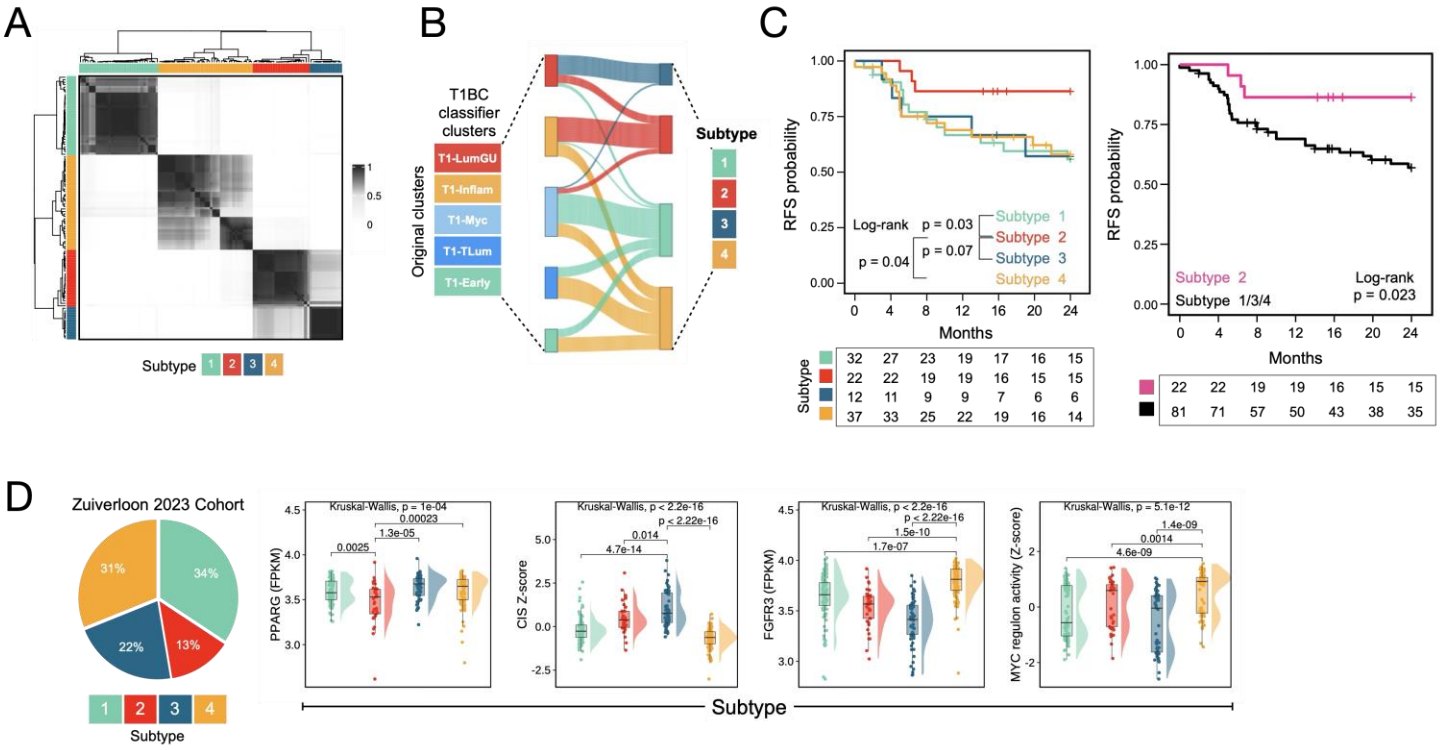


A. Consensus matrix for the four classes, illustrating the level of agreement within each group based on the clustering results. Values closer to 1 indicate higher consensus among samples.

B. Sankey plot comparing sample assignment distribution between the previously described five-cluster NMIBC classes and the four-cluster NMIBC class (ReClusters1-4) described here.

C. Left: Kaplan-Meier curve of recurrence-free survival for patients defined by the ReCluster transcriptomic subtypes with a log-rank p-value for the highlighted comparisons. Right: Kaplan-Meier curve of recurrence-free survival for patients comparing ReCluster 2 vs ReCluster 1+3+4 with a log-rank p-value.

D. Left: Classification of tumors within the BRS-Zuiverloon-2023 Cohort using the described ReCluster classifications. Right: Boxplots comparing expression of PPARG mRNA levels, CIS score, FGFR3 mRNA levels, and MYC regulon activity in tumors from the BRS cohort classified by the ReCluster subtypes. (p-values were calculated from a two-sided Kruskal-Wallis test)

**Supplementary Figure 2:**

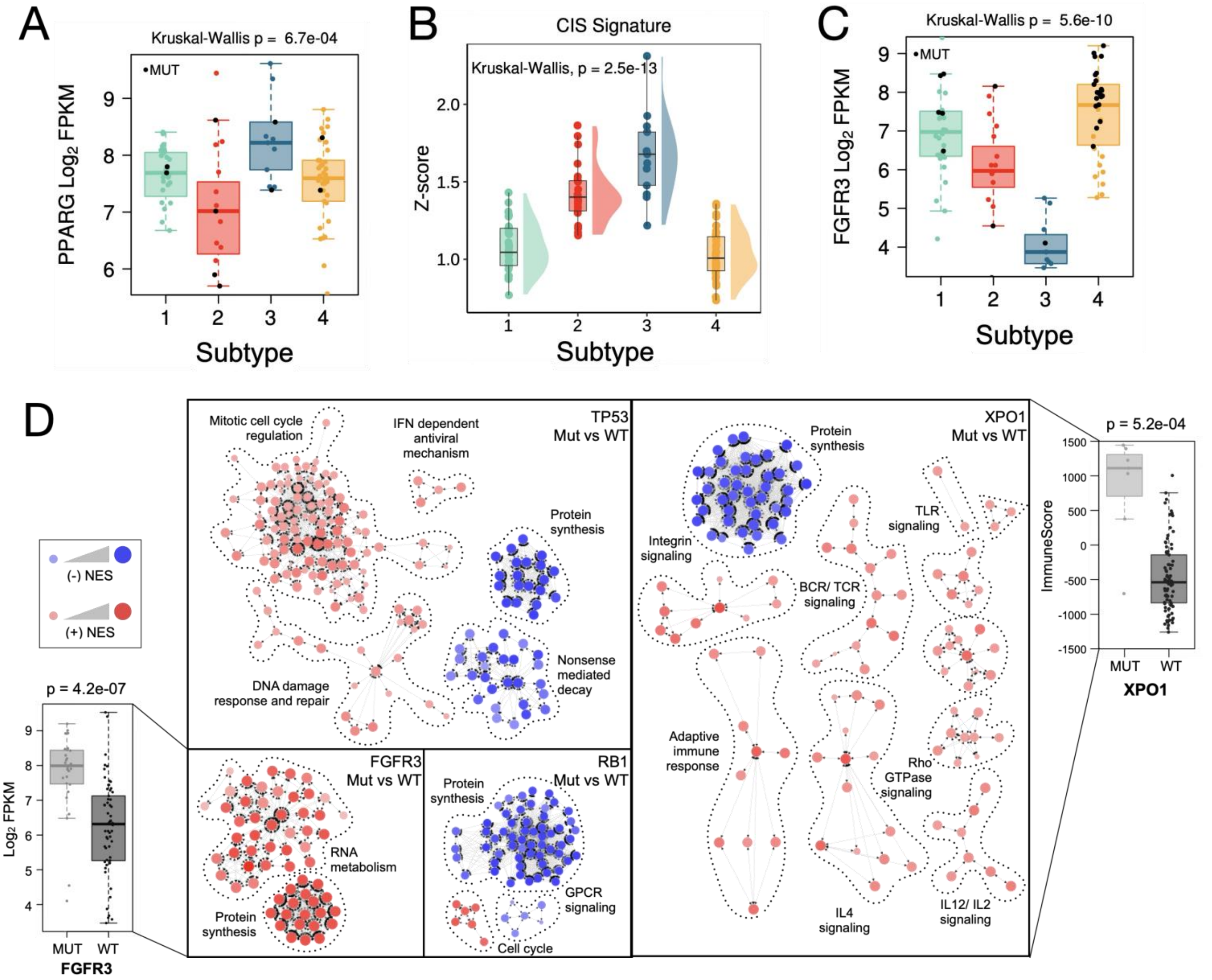


A. Boxplot comparing expression of PPARG mRNA by ReCluster subtypes. (p-value was calculated from a two-sided Kruskal-Wallis test)

B. Plot comparing Z-scored expression of CIS score signature across ReCluster subtypes. (p-value was calculated from a two-sided Kruskal-Wallis test)

C. Boxplot comparing expression of FGFR3 mRNA by ReCluster subtypes. (p-value was calculated from a two-sided Kruskal-Wallis test)

D. Cytoscape network showing gene programs in the highlighted comparisons: Top left – TP53 mutated tumors vs TP53 wildtype tumors, Bottom left – FGFR3 mutated tumors vs FGFR3 wildtype tumors, Bottom right – RB1 mutated tumors vs RB1 wildtype tumors, Right – XPO1 mutated tumors vs XPO1 wildtype tumors.

**Supplementary Figure 3:**

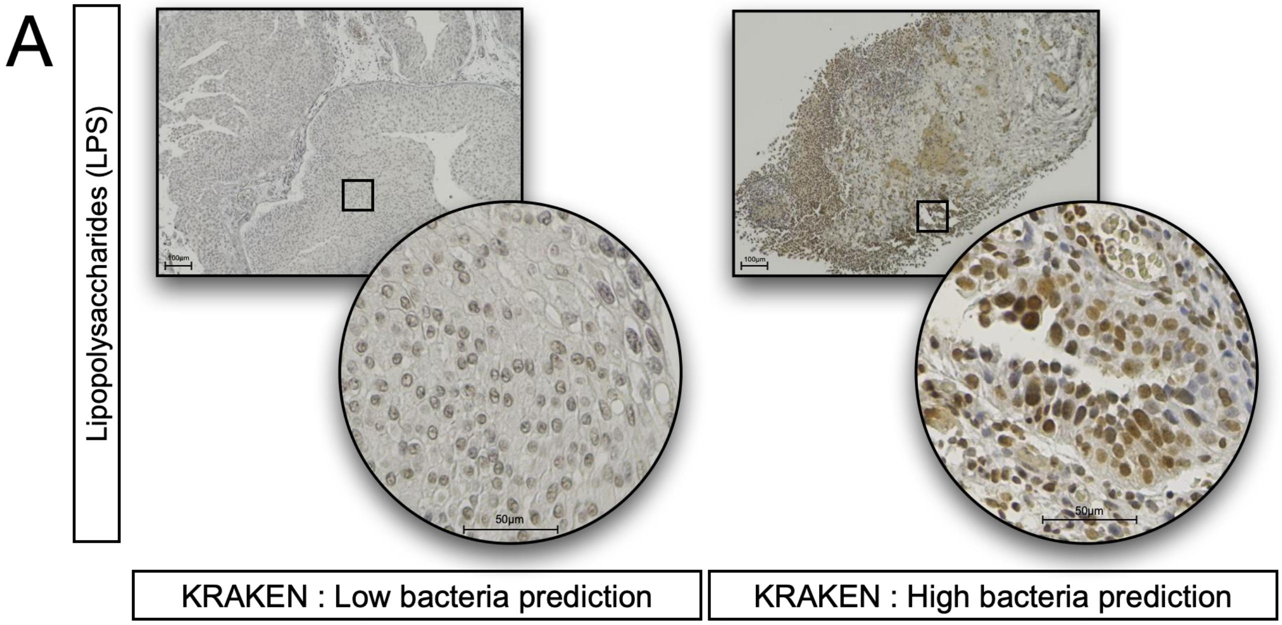


A. Representative immunohistochemistry images showing bacterial lipopolysaccharides (LPS) staining in tumors classified by phylogenetic analysis as having low bacterial reads (top) or high bacterial reads (bottom).
